# Spatial mapping of cell-surface protein glycosylation at molecular resolution

**DOI:** 10.1101/2025.09.30.679542

**Authors:** Dijo Moonnukandathil Joseph, Nathan Linus Sison, Nazlican Yurekli, Sheston Culpepper, Leonhard Möckl

## Abstract

Cell-surface protein glycosylation is a fundamental post-translational modification that plays a crucial role in membrane protein function and cellular behavior. However, elucidating the molecular spatial organization of the cell-surface glycoproteome within the native cellular context has remained challenging due to its structural complexity and density. Here, we introduce a super-resolution microscopy-based approach that enables the spatial analysis of a protein and its glycosylation pattern directly in the cellular environment. Our strategy relies on a combination of lectin-based labeling, metabolic oligosaccharide engineering, and immunocytochemistry. This allows for the mapping of individual protein glycoforms as well as sialylation states of single proteins. By providing access to the spatial axis of the cell-surface glycoproteome, our technique opens up new avenues for understanding the molecular architecture of protein glycosylation in biological processes of central relevance, including, but not limited to, development, signaling, and immune system regulation.

## Main text

Protein glycosylation, the attachment of glycans to proteins, is a ubiquitous post-translational modification in eukaryotic cells.^1^ Protein glycosylation is found both on intracellular and membrane-resident proteins.^2^ It is critical for cellular behavior and control of protein function. Examples include protein folding, protein structure stability, and protein-protein interactions.^3-5^ Protein glycosylation confers a layer of diversity that substantially expands the functional repertoire encoded by the polypeptide sequence. Especially cell-surface proteins carry a plethora of glycan structures; and virtually all cell-surface proteins are glycosylated, establishing the cell-surface glycoproteome.^6^

Unlike transcription and translation, glycosylation is not template-guided. Rather, glycosylation is the product of an ensemble of factors, ranging from transcription levels of glycosyltransferases over pH levels to substrate availability and metabolic state of the cell.^7,8^ Due to this unique complexity and apparent stochasticity, precise analysis of the cell-surface glycoproteome is associated with specific challenges.^9^

Partially, these challenges were addressed in previous years. Prominently, mass spectrometry-based methods have been used.^10,11^ They deliver bulk information on protein glycosylation at high sensitivity for molecular identity, but do not provide any spatial information. Furthermore, electron microscopy has been used to investigate cell-surface glycosylation.^11^ Electron microscopy offers high spatial resolution; however, it yields merely an electron density map, without information on the molecular species imaged. Moreover, both electron microscopy and mass spectrometry generally do not maintain the biological sample in its native state due to comparably invasive sample preparation protocols. More recently, mass spectrometry imaging has been employed to analyze glycosylation under near-native conditions.^12,13^ However, its limited spatial resolution renders it unsuitable for subcellular or molecular-level investigations. Taken together, these previous approaches, while they generated relevant insights into the cell-surface glycoproteome, could not obtain information on spatial glycoproteome organization at molecular resolution in the native environment.

The relative non-invasiveness of light microscopy would make it an ideal method to study the native cell-surface glycoproteome, but the diffraction limit (approx. 250 nm) prevents access to the molecular scale,^14^ rendering this family of methods not suitable for analysis of glycosylation on individual proteins. Proximity-based fluorescence microscopy techniques like Förster Resonance Energy Transfer (FRET) and Fluorescence Lifetime Imaging Microscopy (FLIM) were used to study glycosylation on cells, however, without cellular context.^15,16^

Fortunately, the limited resolution of conventional light microscopy is overcome by super-resolution microscopy techniques, which have yielded revolutionary insights into biological processes.^17-20^ In particular, DNA points accumulation for imaging in nanoscale topography (DNA-PAINT)^21^ and resolution enhancement via sequential imaging (RESI)^22,23^ have proven to be valuable methodologies.

We recently introduced Glycan Atlassing,^24^ a multiplexed super-resolution microscopy pipeline to analyze the communication between nanoscale glycocalyx architecture and cell state. In a second study, we visualized, for the first time, individual sugar moieties within native cell-surface glycans at Ångström resolution.^25^

Extending these protein-agnostic approaches, we here present a method that enables the analysis of protein glycosylation in the native cellular context at molecular resolution. Employing either lectins or metabolic incorporation of unnatural sugars^26^ to label glycosylation and antibody-based labeling to label proteins of interest, the spatial dimension of single-protein glycosylation is analyzed.

Therefore, the approach presented here provides information on the spatial arrangement of different glycoforms of defined proteins with molecular resolution. Indeed, the spatial resolution obtained enables counting of single sugar residues on molecularly-resolved proteins, which defines the limit of the biological length scale. Thus, the spatial axis of the cell-surface glycoproteome is now accessible for the first time.

### Strategy for simultaneous protein and glycosylation labeling in the native cellular context

We present two approaches for spatial analysis of protein glycosylation with molecular resolution. Both approaches enable the simultaneous visualization of proteins along with their associated glycosylation on the cell surface (Fig. 1A). In both cases, the protein of interest is labeled with an antibody-nanobody complex, conjugated to the DNA-PAINT docking sequence R6.^27^ DNA-PAINT-based techniques are employed throughout to determine locations of proteins and glycans or sugar residues.

**Figure 1:**
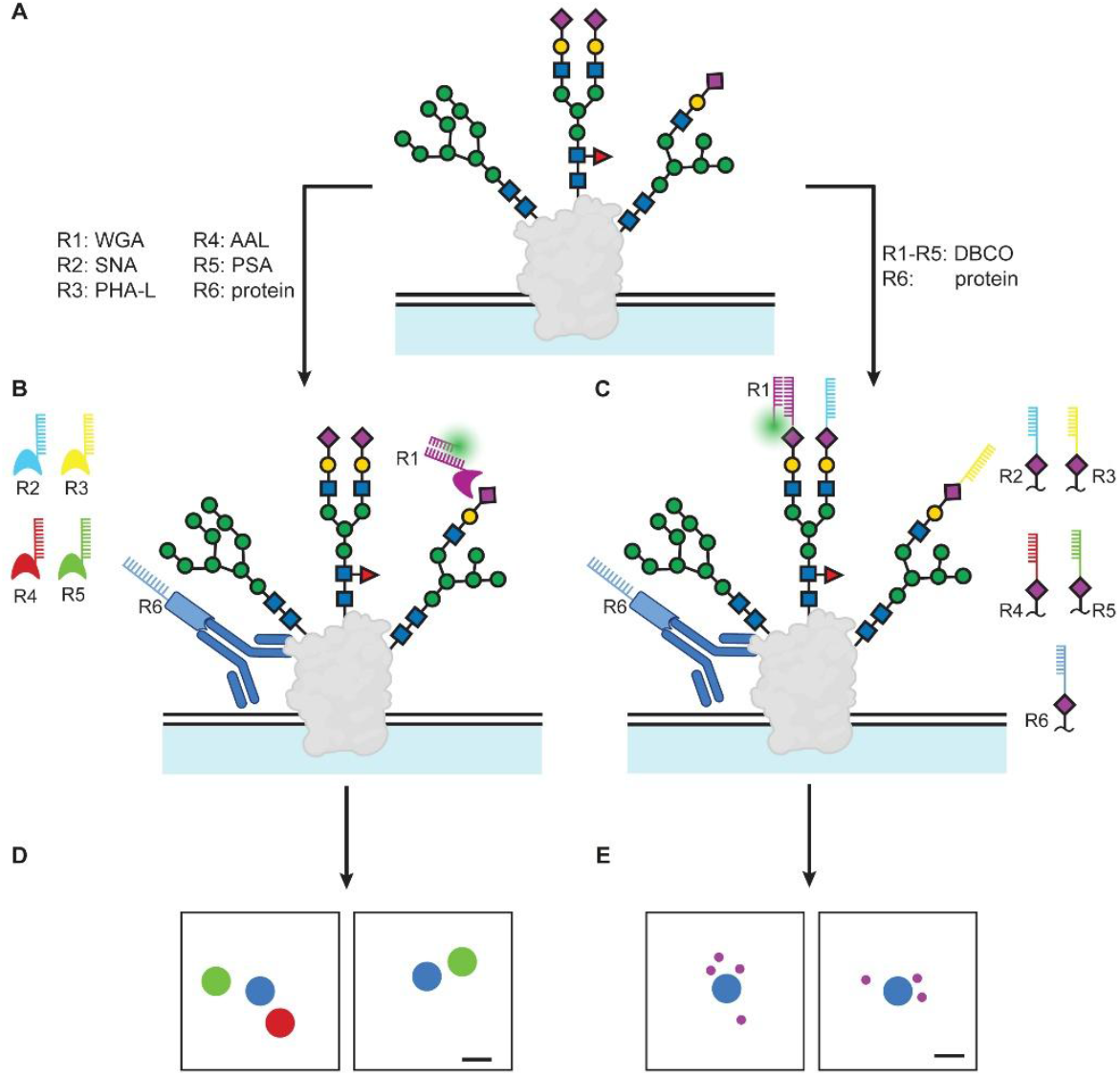
Experimental approach. (A) Schematic depiction of a glycosylated membrane protein. (B) The lectin-based approach relies on the five lectins WGA, SNA, PHAL, AAL and PSA, conjugated to the DNA-PAINT docking sequences R1-R5. (C) The metabolic labeling-based approach relies on metabolic incorporation of N-azidoacetylmannosamine, which enables stochastic labeling of sialic acids with a mixture of the DNA-PAINT docking sequences R1-R5. (D) Protein (blue) and lectin localizations (red/green) are resolved at conventional DNA-PAINT resolution for the lectin-based approach. (E) Protein localizations (blue) are resolved at DNA-PAINT resolution and sugar residues (purple) are resolved at RESI resolution for the metabolic labeling-based approach. Schematics are not to scale. Scale bars in (D) and (E): 15 nm.

For glycosylation readout, the first approach relies on the five lectins WGA, SNA, PHAL, AAL and PSA, conjugated to the DNA-PAINT docking sequences R1-R5, respectively (Fig. 1B). These lectins were chosen based on previous analysis of specificity and affinity as well as labeling scope.^28^ The second approach employs metabolic incorporation of N-azidoacetylmannosamine (Ac_4_ManNAz) to generate azido-functionalized sialic acids,^26^ which are then stochastically labeled with a mixture of the DNA-PAINT docking sequences R1-R5 via live-cell compatible copper-free click chemistry (Fig. 1C).^29^

Together, these approaches provide complementary readouts of protein glycosylation. The lectin-based, multiplexed DNA-PAINT approach yields spatial maps of the protein of interest and five glycan classes, with all six targets resolved at conventional DNA-PAINT resolution (approximately 3 to 5 nm; Fig. 1D). The metabolic labeling-based approach uses conventional DNA-PAINT for the protein of interest combined with RESI for a specific sugar residue, resolving the protein at 3 to 5 nm resolution and the targeted sugar residue at RESI resolution (Fig. 1E).

These two approaches are suited for distinct objectives in spatial mapping of cell-membrane protein glycosylation, and method selection should be guided by the specific research question. When simultaneous profiling of multiple glycan classes is required and a modest reduction in spatial resolution is acceptable, the lectin-based strategy is preferable; for example, to spatially trace organization and redistribution of different protein glycoforms during cellular differentiation. Conversely, when information on a single sugar species with utmost resolution is required, the metabolic labeling approach is preferable; for example, to decipher the role of individual receptor sialylation patterns in immune system regulation and cancer immune escape.

### Multiplexed spatial mapping of protein glycoforms

As outlined above, the lectin-based workflow is grounded in tagging the protein of interest via a DNA-conjugated nanobody-antibody complex and probing the attached glycans with five lectins. At the spatial resolutions achieved here, the labeling footprint – the offset between the true position of the molecular target and the attached label (in this case, the DNA strands) – is non-negligible and must be considered in data analysis. To account for labeling footprint, we used PyMOL-based modeling to estimate the sizes and geometries of the examined proteins and glycans, combining literature information and crystal structure data.^30-33^

Although docking studies can provide plausible orientations of the antibody relative to the target protein, precisely predicting the glycosylation pattern, site occupancy, and steric properties of the glycans is not trivial. Therefore, we adopted a conservative strategy of reporting upper bounds for the labeling footprint and, correspondingly, upper bounds for the maximal separation between protein and glycan labels.

In the configuration illustrated in Fig. 2A, the antibody-nanobody complex extends from the protein in the direction opposite to the glycan of interest. We analyzed three proteins: The epidermal growth factor receptor (EGFR), the glucose transporter 1 (GLUT1), and the matrix metalloprotease 14 (MMP14). In the case of EGFR, the apparent positions of the protein and glycan labels can therefore differ by up to approximately 35 nm. The corresponding upper bounds for GLUT1 and MMP14 are approximately 36 and 37 nm, respectively, reflecting their slightly larger dimensions.

**Figure 2:**
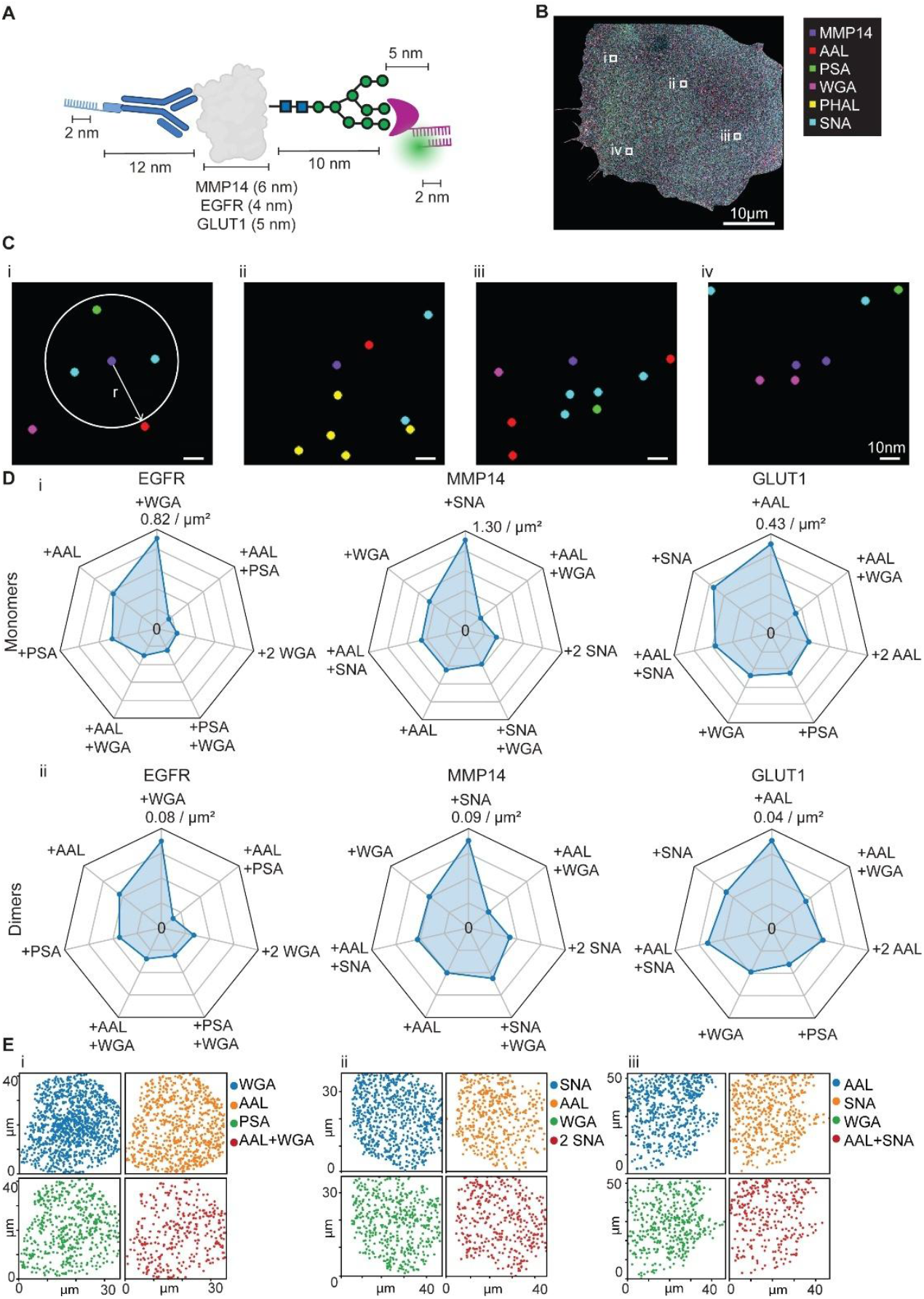
Spatial analysis of EGFR, MMP14, and GLUT1 glycoforms. (A) Estimation of the maximal labeling footprint. Note that the schematic is not to scale. (B) Representative reconstruction of a cell labeled for MMP14 and glycosylation using the five lectins. (C) Zoom-ins corresponding to the areas highlighted in (B). In (i), the search radius based on the maximal labeling footprint estimate is shown. (i-iii) depict MMP14 monomers, (iv) depicts a MMP14 dimer. (D) Distribution of the most prominent glycoforms for each protein of interest. (i) Monomers, (ii) dimers. Lectin occurrence is ordered according to density for monomers. Dimers follow the order of monomers. Averages are shown; n=10 cells for EGFR, n=6 cells for GLUT1, n=6 cells for MMP14. (E) Example images of the spatial distribution of prominent glycoforms for each investigated protein on the cell surface.

Fig. 2B shows a representative multiplexed image from an MCF10AT cell (see Methods and previous work^24^ for details of multiplexed DNA-PAINT acquisition and processing), with MMP14 as the protein of interest. Magnified views of individual MMP14 molecules are provided in Fig. 2C. The four examples illustrate distinct glycosylation patterns observed for MMP14 in its monomeric (i-iii) and dimeric state (iv).

For quantitative analysis, we defined a protein-specific search radius based on the estimated upper bound of the labeling footprint, as described above. Any lectin localization within this radius was assigned to originate from a glycan attached to the corresponding protein. We applied this procedure to each cell independently by iterating over all protein localizations and collecting lectin localizations within the respective search radius. Fig. 2D summarizes the resulting bulk glycosylation readouts for EGFR, GLUT1, and MMP14, averaged across all imaged cells. In Fig. 2E, spatial distribution of four prominent glycoforms of EGFR (i), MMP14 (ii), and GLUT1 (iii) are shown on representative cells.

Strikingly, across the proteins studied, we detected highly distinct glycosylation patterns. A considerable proportion of MMP14 was found to be associated with the lectin SNA, whose putative targets are α(2,6) sialylated glycans. This suggests that on MCF10AT cells, MMP14 monomers predominantly undergo α(2,6) sialylation. Similarly, a significant proportion of GLUT1 were found to undergo core-fucosylation, as evidenced by frequent association of GLUT1 with signals from the fucose-binding lectin AAL. Finally, with respect to EGFR, we predominantly detected N-glycans, with a particular emphasis on N-acetylglucosamine, as suggested by the frequent co-occurence of EGFR localizations with the lectin WGA. These findings are in line with previous literature reports.^34-36^

We would like to emphasize that this constitutes a considerable innovation. Although mass spectrometry can identify glycan compositions and, in favorable cases, site occupancies on a given protein, it requires cell lysis, which discards spatial context and yields ensemble-averaged readouts. In contrast, our workflow assigns glycan class or residue identity to individual proteins on intact cells and localizes them with molecular precision. Consequently, we can determine the location of proteins on the cell surface and its glycosylation state (e.g., α(2,6)-sialylated or core fucosylated, Fig. 2E). We can also quantify the distribution of these glycoforms across nanoscale membrane domains. This capability reveals the spatial heterogeneity and co-organization of glycoforms inaccessible to previous approaches. It enables the molecule-by-molecule mapping of protein glycosylation in the native cellular environment.

All three proteins investigated occur as monomers and dimers on the plasma membrane. Therefore, we searched for spatial signatures of dimerization in the protein localizations. Based on labeling footprint and structural considerations, we identified conservative upper bounds for the protein-protein co-occurrence radii, yielding 36 nm for EGFR, 38 nm for GLUT1, and 40 nm for MMP14. Using these thresholds, we detected dimers for all three proteins, whereas higher-order oligomers – which are biologically not expected – were infrequent.

Next, we analyzed dimer glycosylation using the spatial framework described above. Fig. 2D (ii) shows the resulting glycosylation patterns averaged across all imaged cells. Consistent with the view that none of these proteins is predominantly dimeric in the native state, dimers are less abundant than monomers, as reflected by the lower spatial density of protein localizations. Overall dimer glycosylation resembles that of the corresponding monomers; however, subtle shifts are evident: The most frequent glycosylation class is preserved, but the distribution among the less frequent classes changes. These differences suggest that glycosylation may be involved in modulation of dimerization for the examined proteins – a previously proposed mechanism that, to our knowledge, has not been directly resolved at the single-protein level.^3^

In addition, for EGFR, previous studies suggest prominent N glycosylation,^37,38^ consistent with our data. Some reports suggest that sialylation and fucosylation inhibit EGFR dimerization.^39^ Interestingly, in our analysis, we do not detect high levels of the α(2,6)sialylation-specific lectin SNA on neither monomers nor dimers. Given that we analyzed a transformed cell line with the oncogene KRAS^G12V^ constitutively active,^40^ this might reflect a tendency towards EGFR dimerization, promoting cellular growth.

### Spatial glycoproteomics at single-protein and single-sugar resolution

Next, we analyzed protein sialylation at RESI resolution for individual sugar residues, focusing on EGFR. Similar to the lectin-based analysis, we first determined an upper limit for the labeling footprint and the maximal offset between the protein and sugar labels (Fig. 3A). This yielded a search radius of 30 nm (Fig. 3B).

**Figure 3:**
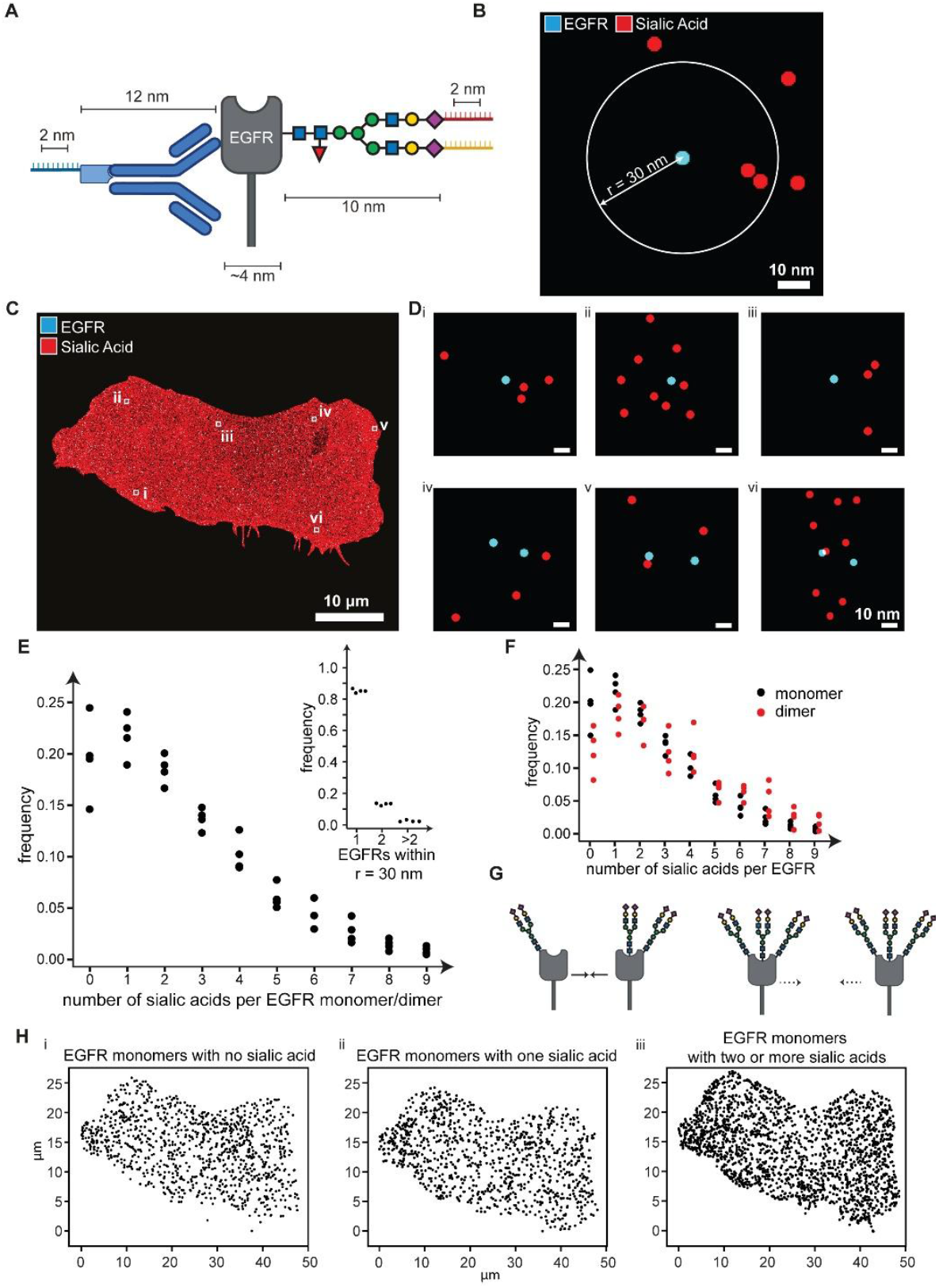
Spatial mapping of EGFR sialylation at single-protein and single-sugar resolution. (A) Estimation of the maximal labeling footprint. Note that the schematic is not to scale. (B) Depiction of the search radius. (C) Representative reconstruction of a cell. EGFR localizations (obtained via DNA-PAINT) are shown in cyan and sialic acid localization (obtained via RESI) are shown in red. Due to the much higher density of sialic acids, EGFR localizations are not visible at this magnification. (D) Representative zoom-ins on individual EGFR localizations. (i-iii) Monomers; (iv-vi) dimers. (E) Number of sialic acid residues around EGFRs. Inset: Number of EGFR monomers, dimers, and higher-order oligomers. n=4 cells. (F) Number of sialic acids around EGFR monomers (black) and dimers (red). n=4 cells. (G) Model for the influence of EGFR glycosylation on EGFR dimerization tendency. (H) Example images of the spatial distribution of EGFR monomers with different sialylation states.

Fig. 3C shows a representative MCF10A cell with EGFR localizations in cyan and sialic acid localizations in red. Because sialic acids commonly cap N-glycans, their density exceeds that of EGFR by a large margin. The zoom-ins in Fig. 3D highlights representative EGFR monomers and dimers, as well as individual sialic acid localizations in close proximity to the protein localizations.

Using the above-described analysis pipeline, we counted individual sialic acid localizations within 30 nm of each EGFR localization without separating monomers and dimers at this stage. Consistent with prior reports of extensive N-glycosylation and sialylation of EGFR,^36^ we found that most EGFR molecules carry at least one, and typically multiple, sialic acids (Fig. 3E). Counts of three or more sialic acids per EGFR were frequent, consistent with the presence of multiantennary, highly sialylated N-glycans or, alternatively, multiple N-glycans, each bearing one or two terminal sialic acids. Notably, sialylation levels were highly consistent across cells. Using the dimer-identification strategy described above, we detected EGFR dimers (Fig. 3E, inset). Reported dimer fractions on cultured cells vary widely, though most studies indicate 5 to 25% of EGFR being dimerized.^41^ Our data align with these estimates, yielding approximately 15% dimers. Higher-order oligomers were rare, supporting the robustness and specificity of our assignments under baseline conditions.

Interestingly, the distribution of sialic acid counts around EGFR monomers and dimers was similar (Fig. 3F). Therefore, when normalized per EGFR subunit, dimers exhibited fewer sialic acids than monomers. This pattern is consistent with a model in which increased sialylation and/or a higher N-glycan load reduces the propensity for dimerization (Fig. 3G). Future work is needed to elucidate the molecular mechanisms underlying this regulatory axis. Finally, we can map the spatial organization of EGFRs based on their different sialylation states at molecular resolution (Fig. 3H), providing a map of EGFR location on the cell surface based on the individual sialylation profile. In summary, the approach presented here is the first to enable direct mapping of EGFR nanoscale organization with simultaneous single-sugar readout of sialylation.

### Conclusion

This study introduces a comprehensive approach for the simultaneous imaging of proteins and their associated glycosylation. By combining antibody-based labeling of proteins with either lectins or metabolic incorporation of unnatural sugars to label glycosylation, we successfully mapped protein glycosylation in the native biological state at single-protein and single-glycan/ -sugar resolution, a level of detail that was previously unattained.

Glycosylation is one of the most complex posttranslational modifications, with a range of biological processes contributing. A functional interplay between metabolic state, pathological cellular events, protein glycosylation, and cellular behavior seems conceivable. Findings to date, however, could not yet be integrated into a comprehensive model. The approach introduced here enables the direct analysis of spatial protein glycoform organization. This will enable to delineate the functional involvement of different protein glycosylation states for membrane architecture and cellular behavior.

The strategy we present is rooted in the tantamount biological relevance of cell-surface glycosylation. One approach investigates protein glycosylation with a focus on the structural diversity of glycans, employing lectins targeted at prominent structural motifs in conjunction with multiplexed imaging. The second approach resolves single sugar residues (in our case, sialic acids) within glycans on individual proteins.

For example, cancer cell sialylation has been proven to directly modulate immune cell activity.^42,43^ This has led to the development of cancer-targeted therapeutics that directly cleave cancer cell sialylation in order to improve patient prognosis.^44^ The approach presented here directly allows for quantification of immune and cancer receptor sialylation. This will enable studies of glycan-mediated immune evasion as well as its therapeutic reversal with a previously not attainable scope.

While our strategy provides a new perspective on cell-surface glycoproteome analysis, it still has limitations. First, we focus analysis to a single protein of interest here. Analyzing the relevance of protein glycosylation on protein-protein interactions will be a highly useful expansion of the principle presented here. In order to achieve this, an expansion of the labeling toolbox is required, as currently only six speed-optimized DNA-PAINT sequences exist.^27^ However, employing adapter strands (FLASH-PAINT)^45^ or recycling of sequences (SUM-PAINT)^46^ is possible and should enable such investigations. Second, we do not analyze protein localization at RESI resolution. Using adapter strands or sequence recycling, it would be possible to perform RESI on both the protein and the glycosylation pattern. However, it remains to be investigated if the resolution gain is universally required as conventional DNA-PAINT resolution should, at least for monomeric proteins expressed at typical levels, suffice to separate individual copies. Third our method currently employs antibody-based labeling of the protein, which introduces a considerable labeling footprint. More targeted methods like unnatural amino acid incorporation could mitigate the labeling footprint. However, they would require genetic engineering of native proteins. Finally, our approach currently investigates a maximum of five glycan (sub-)structures. A larger number would yield a more comprehensive picture, similar to the one obtained in mass spectrometry-based analyses. We would like to point out, however, that this limitation is mainly attributable to the scarcity of labeling agents that can specifically and selectively target glycans. Advances in this area will substantially improve the scope of the method presented here.

In summary, our strategy enables the precise analysis of protein glycosylation with single-protein and single-sugar resolution. We therefore provide the field of biology with an approach to study cell-surface protein organization from a fundamentally new perspective. In addition, considering the emerging relevance of glycobiology in clinical and translational research as well as precision medicine,^42,47,48^ we believe that the ability to spatially map individual glycoforms will be valuable in the development of diagnostic and therapeutic targets.

## Acknowledgements

The authors gratefully acknowledge financial support from the Else-Kröner-Fresenius-Stiftung (grant ID 2020_EKEA.91 to LM), the German Research Foundation (DFG, grant ID 529257351 to LM), the Wilhelm-Sander-Stiftung (grant ID 2023.025.1 to LM), and the Max Planck Society. Helpful discussions with and scientific input from Karim Almahayni and Nikhitha Murali Shankar are gratefully acknowledged.

## Materials and Methods

### Cell culture

MCF10A/AT cells were cultured in T-25 flasks (Greiner Bio-one, cat# 690175) in culture medium containing DMEM/F12 (Gibco, ref# 21041-025) and supplemented with 5% horse serum (Gibco, ref# 16050-112), 20 ng/mL epidermal growth factor (EGF) (Gibco, ref# PHG0311), 0.5 μg/mL hydrocortisone (Sigma-Aldrich, ref# H0396), 100 ng/mL cholera toxin (Sigma-Aldrich, cat# C8052-5MG), 10 μg/mL insulin (Sigma-Aldrich, cat# I1882-100MG), and 1% penicillin/streptomycin (Sigma-Aldrich, cat# P0781-100ML). Cells were incubated at 37°C in a humidified atmosphere with 5% CO_2_. After reaching 70-80% confluency, cells were washed with 5 ml of DPBS, Ca^2+^/Mg^2+^-free (Gibco, ref# 14190-094) and split by brief incubation with 0.05% trypsin-EDTA solution (Gibco, ref# 2500-054) for 8-10 minutes. Following trypsin neutralization with medium including FBS, cells were centrifuged at 3000 rpm for 5 minutes, and the resulting pellet was resuspended.

### Cell seeding and fixation

Cultured cells were seeded into 8-well chamber slides (Ibidi GmbH, ref# 80807-90) and incubated for 24 hours for prior sample preparation. Samples were washed with DPBS, Ca^2+^/Mg^2+^ (Gibco, ref# 14040-091) 3 times. Then 4% paraformaldehyde, diluted with DPBS, Ca^2+^/Mg^2+^-free, from a 16% stock solution (Thermo Scientific, ref.no: 28908) was added to wells and cells were incubated at room temperature for 15 minutes. Then, cells were washed three times with DPBS, Ca^2+^/Mg^2+^-free.

### Metabolic incorporation of unnatural azido sugars

After 6 hours of incubation of cells after seeding, culture medium was exchanged with fresh culture medium supplemented with 50 μM Ac_4_ManNAz (Thermo Fisher Scientific) and cells were incubated for 48 h. After incubation, copper-free click chemistry was performed. Cells were washed 3 times with DPBS Ca^2+^/Mg^2+^. A solution containing 50 μM of each of the 5 docking strands conjugated to DBCO (sequence R1-R5) were prepared in complete culture medium. Cells were treated with this solution for 2 h at 37 °C. Following this incubation, the samples were washed with DPBS, Ca^2+^/Mg^2+^ (Gibco, ref# 14040-091) 3 times. Then, cells were fixed as described above. After this, cells were washed 3 times with DPBS, Ca^2+^/Mg^2+^-free. The cells were then permeabilized with 0.1% Triton-X (Alfa Aesar, cat# A16046) in DPBS, Ca^2+^/Mg^2+^-free for 10 min at room temperature. Finally, cells were washed 3 times with DPBS Ca^2+^/Mg^2+^-free.

### Immunocytochemistry

The desired primary antibody (Abcam anti-EGFR antibody EP38Y, ab52894; anti-MMP14 antibody EP1264Y, ab51074; Abcam anti-GLUT1 antibody EPR3915, ab115730) was pre-fused with the nanobody (Massive Photonics Donkey anti-rabbit IgG 5xR6) in the antibody incubation buffer for 2 h. The antibody incubation buffer was prepared in DPBS, Ca^2+^/Mg^2+^-free containing 1 mM EDTA, 0.02% Tween-20, 0.05 mg/mL salmon sperm DNA (Invitrogen, cat#15632011), 0.05% sodium azide, and 2% (w/v) BSA (Sigma-Aldrich, cat# A9647-100G). Components were added sequentially to PBS, mixed until fully dissolved, and the final volume was adjusted with DPBS, Ca^2+^/Mg^2+^-free.

Then, an excess (molar ratio of 1:2) of unlabeled blocking nanobody was added to the mixture for 5 min (NanoTag Biotechnologies, cat: K0102-50). The addition of the unlabeled secondary nanobody should take place just before the prefusion mixture is added to the sample to reduce the chances of the unlabeled nanobody sticking to a non-specific target.

After permeabilization, the cells were incubated in a blocking buffer for 1 h at room temperature. The blocking buffer was prepared in DPBS, Ca^2+^/Mg^2+^-free containing 0.05 mg/mL salmon sperm DNA, 0.02% Tween-20, and 3% (w/v) BSA. All components were dissolved in PBS and the volume adjusted accordingly. Then, the cells were washed three times with DPBS, Ca^2+^/Mg^2+^-free. The pre-fusion mixture was then added to the cells and allowed to incubate for 2 h at room temperature. After that, the cells were washed five times with DPBS, Ca^2+^/Mg^2+^-free at room temperature. In each washing round, the cells were placed on a shaker rotating at 50 RPM for five minutes.

### Lectin labeling

After fixation, cells were permeabilized with 0.1% Triton-X (Alfa Aesar, cat# A16046) for 10 minutes at room temperature, followed by four DPBS, Ca^2+^/Mg^2+^-free washing steps. A lectin cocktail (2.5 μg/mL of each lectin, conjugated to DNA strands R1 to R5 by Massive Photonics) prepared in 1X tris buffer (Fisher Bioreagents, ref# M-15836) was applied to the cells at room temperature for 30 min, allowing specific binding to glycan targets on the cellular surface. Following incubation, cells were washed three times with DPBS, Ca^2+^/Mg^2+^-free. Prior to imaging, another permeabilization step using 0.1% Triton-X 100 was performed for 10 min at room temperature, followed by four washing steps with DPBS, Ca^2+^/Mg^2+^-free

### Optical setup

Fluorescence imaging was carried out on an inverted microscope (Nikon Instruments, Eclipse Ti2) with the Perfect Focus System. Objective-based Total Internal Reflection Fluorescence (TIRF) mode was used, employing a high NA objective (Nikon Instruments, Apo SR TIRF×100, NA 1.49, Oil) and the Nikon manual TIRF module. Excitation was done using a 560-nm laser (MPB Communications, 1 W) coupled into the microscope via the TIRF module. Excitation power was controlled in free space using a filter wheel (Thorlabs, FW212CNEB). The beam was cleaned using a filter (Chroma Technology, ZET561/10) and coupled into the microscope objective using a beam splitter (Chroma Technology, ZT561RDC). Fluorescence emission was spectrally filtered using an emission filter (Chroma Technology, ET600/50m, and ET575LP) and imaged on an sCMOS camera (Hamamatsu Orca Fusion) without further magnification, resulting in an effective pixel size of 130 nm after 2×2 binning. TIRF illumination was used for all measurements with a laser power of approx. 33 mW above the objective corresponding to an intensity of ∼170 W cm-2. The central 1152×1152 pixels (576×576 after binning) of the camera were used as the region of interest. Raw microscopy data was acquired using μManager (Version 2.0.3).

### Sequential DNA-PAINT

Fixed samples prepared for imaging were incubated with gold nanoparticles (Cytodiagnostics, #G-90-100) at a 1:3 dilution in MilliQ^®^ water for 10 min. After incubation, the sample was washed 5 times with DPBS, Ca^2+^/Mg^2+^-free to remove excess nanoparticles.

Imaging buffer was prepared by combining 50 ml of DPBS, Ca2+/Mg2+-free with 0.0146 g ethylenediaminetetraacetic acid (EDTA) (PanReac, cat# 60-00-4), 1.461 g sodium chloride (NaCl) (Alfa Aesar, cat# A12313), and 10 µl TWEEN-20 (MP Biomedicals, cat# 103368) in a 50-ml falcon tube. All components were thoroughly mixed until completely dissolved. The prepared buffer was stored at 4°C until use in subsequent imager strand preparation. Imaging strands for DNA-PAINT were prepared by diluting 1 µM stock solutions in the buffer.

For the RESI approach, imaging strands are diluted with the imaging buffer to the following concentrations: R1, R2 and R4: 100 pM, R3: 12.5 pM, R5: 150pM, and R6: 250pM. For each RESI channel data, as well for the protein (R6), 40,000 frames with an exposure time of 100 ms were acquired.

For the lectin approach, imaging strands are diluted with the imaging buffer to the following concentrations: R1, R3, R5 and R6: 250 pM, R2: 75 pM, and R4: 125pM. For each lectin, as well as for the protein (R6), 20,000 frames were acquired with 100 ms exposure time.

For both approaches, six subsequent imaging rounds were performed. Between each round, thorough washing was performed to remove the imager strands from the previous imaging round. This was observed in real time on the microscope.

### Analysis of raw data and postprocessing

Raw fluorescence data was reconstructed using the Picasso software package, version 0.8.^49^ An intensity threshold of 5000 was used to detect the localization events in each frame. An estimate of experimental localization precision was calculated using Nearest Neighbor Analysis (NeNA precision).^50^ For the Lectin approach, Redundant cross correlation (RCC) followed by fiducial based correction was used for drift correction in each imaging round. For the metabolic labeling approach, RCC followed by Adaptive Intersection Maximization (AIM) method was used for drift correction. This route was chosen due to the higher number of localizations in this case.^51^ For each RESI imaging channel, the following AIM parameters were applied: intersection distance: 3xNeNA precision and maximum drift in segment: 3x intersection distance.

Data from six imaging targets were aligned using the localized position of all gold nanoparticles as fiducial markers throughout each channel. After alignment, the areas of interest (single cells) were segmented. We used a modified version of the Picasso software package in python^52^ for all analysis after segmentation until clustering. Repeated sampling of a docking site by transient DNA-DNA interactions was clustered into a single docking site location using a radius of 2.35 times the experimental NeNA precision. In the lectin dataset, we achieved a sub-10 nm NeNA precision, for RESI data from labelled sialic acids, average NeNA from 5 channels (3.33 nm) were used. For the lectin approach, a minimum of two sampling events were set as a threshold for a cluster, with further strict filtering for interaction kinetics. For the metabolic approach, a minimum of five sampling events were set for a threshold. For each cluster, duration of each DNA-DNA interaction was analyzed to account for unspecific sticking events. If a time bin of 200 frames (1% of the total length of the stack) in a single cluster contains more than 90% of all the events in that cluster, the respective cluster is rejected. Thus, single and/or atypically extended events are rejected as false localization. Cluster centers are calculated and are used as the location of glycan targets.

### Spatial analysis

All spatial analyses are performed at the level of single cells on cluster centers. If two proteins were found closer than a distance corresponding to their labeling footprint (protein plus antibody complex plus docking strand), they were defined as dimers.

For monomeric protein glycosylation analysis, a search radius depending on the size of protein and considering labeling footprint were defined (length of DNA strands, antibody-nanobody complex and lectin). For each protein location, a search is conducted for associated lectin/sialic acid locations. These detected lectin/sialic acid locations across all the lectin channels are grouped together with the corresponding protein to form the monomer protein glycosylation. This “search and group” was repeated for all protein locations and the density of protein glycoforms per cell is estimated. For the lectin-based analysis the spatial density distribution of seven predominant glycoforms are plotted in a spider plot, and the locations of top 5 glycoforms on the cell are mapped.

For dimeric protein glycosylation analysis, a search radius for dimers considering the labeling footprint as before is defined for each protein. The location of individual proteins in the dimers are stored. Then for each protein in the dimer, a search for glycosylation is performed as in the monomeric case, and the detected glycosylation for each monomer is coalesced to generate the respective dimer glycoform. For the lectin-based analysis, the spatial density distribution of seven predominant dimer glycoforms are plotted in a spider plot.

